# Potent Neutralization of Omicron and other SARS-CoV-2 Variants of Concern by Biparatopic Human VH Domains

**DOI:** 10.1101/2022.02.18.481058

**Authors:** Chuan Chen, James W. Saville, Michelle M. Marti, Alexandra Schäfer, Mary Hongying Cheng, Dhiraj Mannar, Xing Zhu, Alison M. Berezuk, Anupam Banerjee, Michele D. Sobolewski, Andrew Kim, Benjamin R. Treat, Priscila Mayrelle Da Silva Castanha, Nathan Enick, Kevin D McCormick, Xianglei Liu, Cynthia Adams, Margaret Grace Hines, Zehua Sun, Weizao Chen, Jana L. Jacobs, Simon M. Barratt-Boyes, John W. Mellors, Ralph S. Baric, Ivet Bahar, Dimiter S. Dimitrov, Sriram Subramaniam, David R. Martinez, Wei Li

## Abstract

The emergence of SARS-CoV-2 variants of concern (VOCs) requires the development of next-generation biologics that are effective against a variety of strains of the virus. Herein, we characterize a human V_H_ domain, F6, which we generated by sequentially panning large phage displayed V_H_ libraries against receptor binding domains (RBDs) containing VOC mutations. Cryo-EM analyses reveal that F6 has a unique binding mode that spans a broad surface of the RBD and involves the antibody framework region. Attachment of an Fc region to a fusion of F6 and ab8, a previously characterized V_H_ domain, resulted in a construct (F6-ab8-Fc) that neutralized Omicron pseudoviruses with a half-maximal neutralizing concentration (IC_50_) of 4.8 nM *in vitro*. Additionally, prophylactic treatment using F6-ab8-Fc reduced live Beta (B.1.351) variant viral titers in the lungs of a mouse model. Our results provide a new potential therapeutic against SARS-CoV-2 VOCs - including the recently emerged Omicron variant - and highlight a vulnerable epitope within the spike protein RBD that may be exploited to achieve broad protection against circulating variants.

## Introduction

Since the outbreak of the coronavirus disease 2019 (COVID-19) pandemic [1–3], more than 415 million cases and 5.83 million deaths have been confirmed as of February 16^th^, 2022. To treat infection by severe acute respiratory syndrome coronavirus 2 (SARS-CoV-2), the causative agent of COVID-19, various therapeutics have been explored, such as convalescent patient sera [4], neutralizing antibodies (nAbs) [5–15], and small antiviral molecules [16–21]. The spike glycoprotein (S protein), which engages the human angiotensin converting enzyme 2 (hACE2) receptor [22], is a major target for Ab-mediated neutralization. nAbs that block S protein - ACE2 binding are promising therapeutic candidates, with a plethora of SARS-CoV-2 nAbs reported - several of which have received emergency use approval (EUA) in the US [9, 23–26].

The receptor binding domain (RBD) within the S1 region of the S protein exhibits a high degree of mutational plasticity and is prone to accumulate mutations that lead to partial immune-escape [27–33]. The World Health Organization (WHO) has designated several SARS-CoV-2 lineages as Variants of Concern (VOCs), which are more transmissible, more pathogenic, and/or can partially evade host immunity, including the Alpha, Beta, Gamma, Delta variants and the recently identified Omicron variant [27, 34–39]. Some pan-sarbecovirus mAbs have been demonstrated to retain their neutralization activity against these VOCs [40, 41]. The Omicron variant is heavily mutated and contains 30 amino acid changes in its S protein, with 15 of mutations localizing to the RBD [42]. Some of these mutations have been predicted or demonstrated to either enhance transmissibility [43] or to contribute to escape from most nAbs that were raised against the original (Wuhan-Hu-1) or early VOCs lineages of SARS-CoV-2. The continuous evolution and emergence of VOCs that can partially evade host immunity requires the development of Abs with broad neutralizing activity that can block or reduce disease burden. Additionally, multi-specific Abs or Ab cocktails hold promise to resist mutational escape by targeting multiple epitopes on the SARS-CoV-2 S protein [15, 39]. Recently, several bispecific Abs have been reported that show broad neutralization efficacy against SARS-CoV-2 variants [6, 44–46], therefore the generation of bispecific or multi-specific nAbs to target variants that otherwise evade immune response is a viable therapeutic strategy.

In this study, we identify a V_H_ domain (V_H_ F6) which shows broad neutralizing activity against SARS-CoV-2 VOCs including the Alpha, Beta, Gamma, Delta, and Omicron VOCs. V_H_ F6 binds a relatively conserved portion of the receptor binding motif (RBM), using a unique framework region (FR)-driven paratope. By combining V_H_ F6 with our previously identified Ab, V_H_ ab8, we developed a biparatopic Ab (F6-ab8-Fc), which exhibits potent neutralizing activity against all SARS-CoV-2 VOC psuedoviruses (including the Omicron variant) and several live virus variants. Prophylactic dosing with F6-ab8-Fc also reduced viral titres in the lungs of a mouse model and high doses of F6-ab8-Fc protected against mortality. Our study identifies a novel broadly neutralizing V_H_ domain Ab with a unique paratope and provides a potent biparatopic Ab (F6-ab8-Fc) against all SARS-CoV-2 VOCs, including the presently dominant Omicron variant.

## Results

### Identification of a novel antibody domain (V_H_ F6) which binds to most prevalent RBD mutants and neutralizes SARS-CoV-2 VOCs

To identify cross-reactive V_H_ domains against SARS-CoV-2 VOCs, we adopted a sequential panning strategy to pan our in-house large V_H_ phage library. We used E484K mutated RBD for the first round of panning, wild type (WT) RBD for the second, and the S protein S1 domain containing K417N, E484K, and N501Y mutations for the third (**Fig. S1A)**. Following these three rounds of panning, a dominant clone, V_H_ F6, was identified by ELISA screening. V_H_ F6 bound to the WT and Beta RBDs with half-maximal binding concentrations (EC_50_) of 5.1 nM and 7.2 nM respectively (**Fig. S1B)**. V_H_ F6 also bound to the WT, Alpha, and Beta S1 proteins (**Fig. S1C)**. To assess the cross-reactivity of V_H_ F6, we performed ELISA and pseudovirus neutralization assays. V_H_ F6 was able to bind to trimeric spike proteins from multiple SARS-CoV-2 VOCs including Alpha, Beta, Gamma, Kappa, and Delta variants **(Fig. S1D)**. Furthermore, we evaluated the ability of V_H_ F6 to bind RBDs containing single-point mutations at mutational sites commonly observed in currently circulating variants. V_H_ F6 bound to 35 out of the 37 assayed RBD mutations, with only F490S and F490L mutants escaping binding (**Fig. 1A and Fig. S1E**). V_H_ F6 was able to neutralize ancestral SARS-CoV-2 (WT), Alpha, Beta, Gamma, and Delta spike pseudotyped viruses with 50% pseudovirus neutralizing Ab titers (IC_50_) of 28.47, 40.32, 3.62, 6.23, and 0.40 nM respectively (**Fig. 1B)**.

**Fig. 1.**
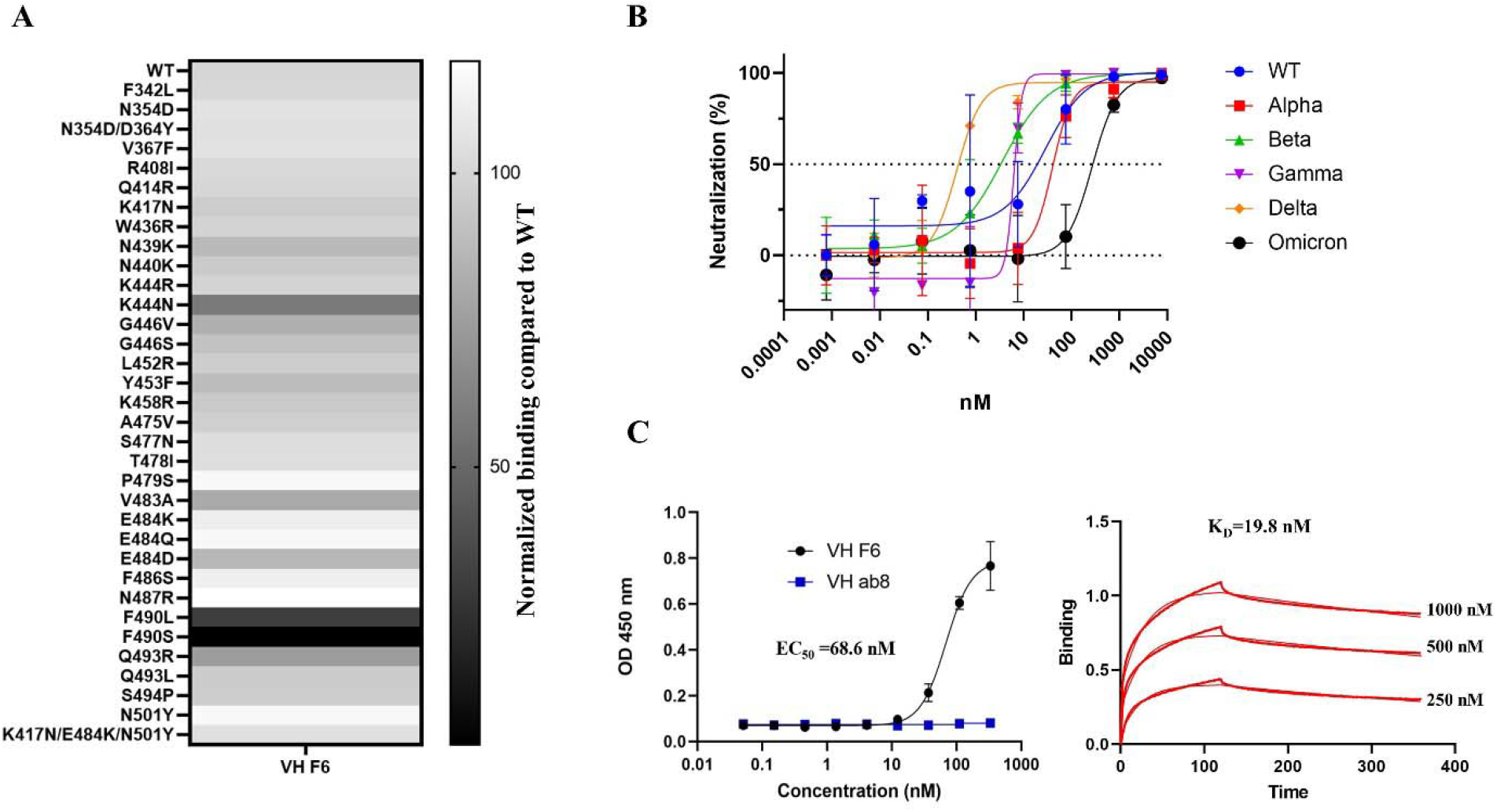
F6 binds to prevalent RBD mutants and neutralizes SARS-CoV-2 VOCs including Omicron. **A.** Heat map of V_H_ F6 binding to circulating RBD mutants. The binding of V_H_ F6 to RBD mutants was detected by ELISA and normalized by comparing area under the curves (AUCs) between mutant and wild type RBD. **B**. Neutralization of SARS-CoV-2 WT, Alpha, Beta, Gamma, Delta, and Omicron variants pseudovirus by V_H_ F6. Experiments were performed in duplicate and error bars denote ± SD, n=2. **C**. Measurement of V_H_ F6 binding to the Omicron RBD by ELISA (left) and BlitZ (right).

The Omicron variant escapes most mAbs that are in clinical use [47]. We found that V_H_ F6 was able to bind the Omicron RBD with an EC_50_ of 68.6 nM as tested by ELISA, which is consistent with the binding dissociation constant (*K*_*D*_= 19.1 nM) obtained by BlitZ (**Fig. 1C**). Furthermore, V_H_ F6 neutralized Omicron psuedoviruses with an IC_50_ of 269 nM (**Fig. 1B**).

### CryoEM structure of the V_H_ F6 - Beta variant spike protein complex reveals a unique FR-driven binding mode

To garner structural insights into the broad neutralization exhibited by V_H_ F6, we solved the cryo-electron microscopy (cryoEM) structure of V_H_ F6 bound to a prefusion stabilized Beta S trimer at a global resolution of 2.8 Å (**Fig. S2 and Table S1**). The Beta variant trimer was chosen for this structural analysis as it contains K417N, E484K, and N501Y mutations, which are partially present in other variants of concern (Alpha, Gamma, and Omicron). The cryoEM reconstruction revealed density for three bound V_H_ F6 molecules with strong density observed for V_H_ F6 binding to a “down” RBD, and moderate or weak densities for two V_H_ F6 molecules binding “up” RBDs (**Fig. 2A**). The strong density for V_H_ F6 bound to the “down” RBD enabled focused refinement, providing a local resolution density map at 3.0 Å and enabling detailed analysis of the V_H_ F6 epitope (**Fig. 2B**).

**Fig. 2.**
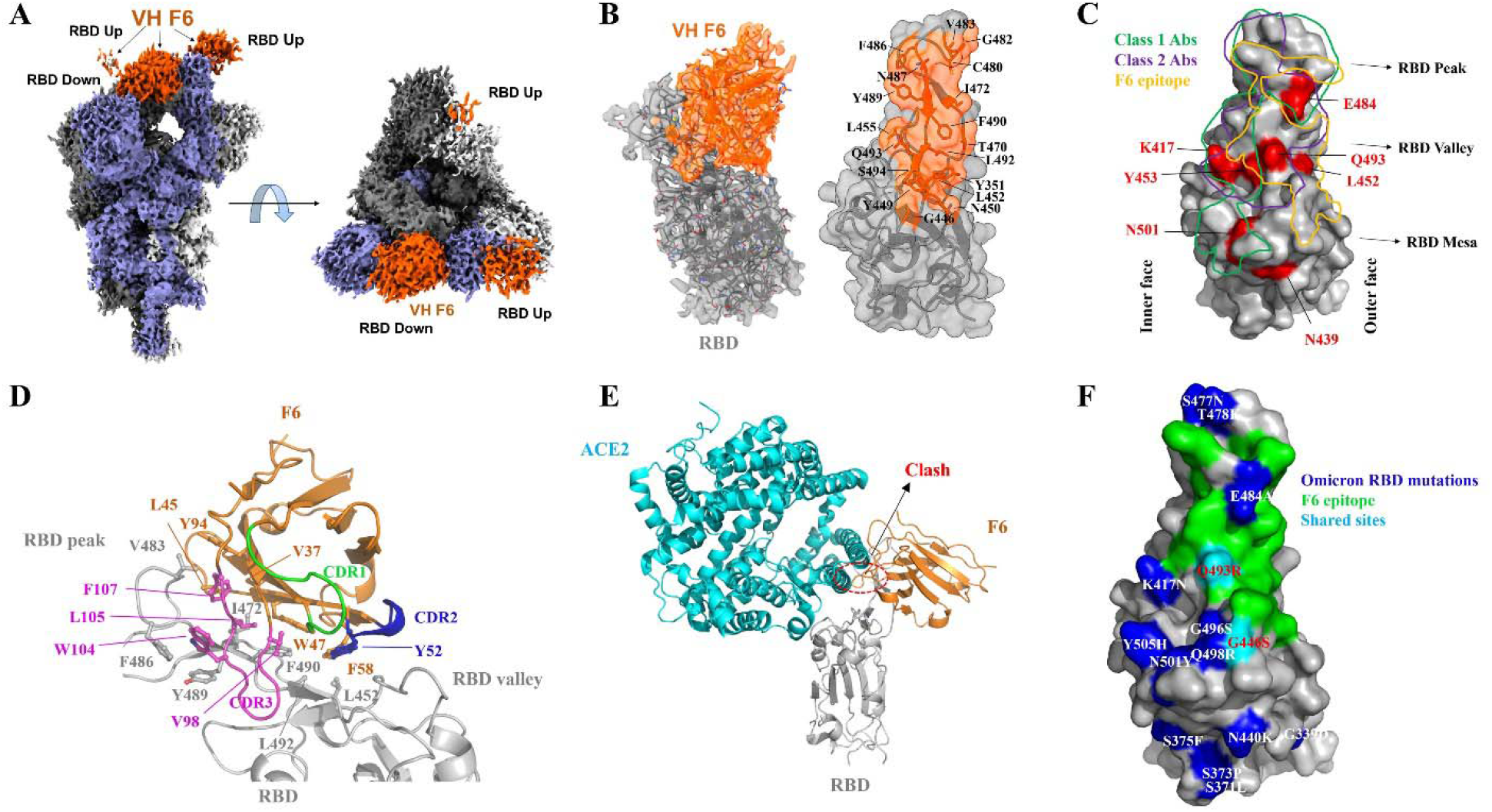
CryoEM structure of V_H_ F6 in complex with the SARS-CoV-2 Beta variant spike protein. **A.** Global cryoEM map of the Beta variant spike protein in complex with V_H_ F6. Density corresponding to the Beta variant trimer is colored in shades of grey and violet while density corresponding to V_H_ F6 molecules is colored in orange. **B**. Left: Focus refined density map of the Beta variant RBD - V_H_ F6 complex with docked atomic model. Right: Molecular surface representation of the epitope of V_H_ F6 on the Beta variant RBD. The side chains of residues within the binding footprint of V_H_ F6 are displayed and colored orange. **C**. Footprints of class 1 Abs (green), class 2 Abs (purple), and V_H_ F6 (orange) on the molecular surface of the SARS-CoV-2 RBD. Commonly mutated and antibody-evading mutations are colored in red. **D**. Focused view of the atomic model at the V_H_ F6 - RBD interface. The side chains of discussed residues are shown, with the scaffold colored orange, CDR1 green, CDR2 blue, CDR3 magenta and the RBD gray. **E**. Superposition of V_H_ F6-RBD (orange) and ACE2-RBD (cyan) complex atomic models. The RBD is shown in grey and the ACE2-RBD model was derived from PBD ID: 6m0j. **G**. Mapping the Omicron mutations onto the RBD structure with comparison to the F6 epitope

V_H_ F6 binding spans the RBD “peak” and “valley” regions, with its footprint skewed towards the RBD “outer face” (**Fig. 2B and 2C**). This interface is exposed in both “up” and “down” RBD conformations, thus explaining V_H_ F6 binding to both states simultaneously. Interestingly, the framework regions (FRs) of F6 – a heavy-chain (V_H_) only Ab – expands the interaction interface beyond the conventional complementarity-determining regions (CDRs) (**Fig. 2D)**. Specifically, the hydrophobic FR2 residues present a hydrophobic core that associates with hydrophobic RBD residues which line the RBD peak and valley regions. This large FR engagement contributes an interaction area that accounts for up to 36% of the total antibody paratope. Such substantial involvement of FRs causes V_H_ F6 to adopt an atypical perpendicular binding angle relative to the RBD, with its FR2, FR3 and CDR3 wrapping around the RBD peak (**Fig. 2D)**. In addition to FRs, CDR2 and CDR3 also contribute to the RBD binding interaction via hydrogen bonding, π-π stacking and van der Waals interactions (**Fig. S3C-E**). Due to its positioning toward the RBD outer edge, the V_H_ F6 footprint only slightly overlaps with the hACE2 binding interface, potentially rationalizing its weaker RBD binding competition with hACE2 as compared to ab8 [48, 49] (**Fig. S3A, S3B and Fig. 2E)**.

The V_H_ F6 bound Beta S protein structure rationalizes the broad activity of V_H_ F6 against various RBD mutants. K417, N501 and E484 – frequently mutated in VOCs and imparting escape from many nAbs – are not within the V_H_ F6 epitope (**Fig. 2C)**. The RBD residue Q493, which is mutated in the Omicron variant and induces escape from the clinical Ab REGN10933 [50], is located within the V_H_ F6 epitope to form hydrogen bonds with the main chain of G101 and S102 in the CDR3 (**Fig. S3C)**. Despite these specific hydrogen bonds, the Q493R/L mutations did not significantly impact V_H_ F6 binding (**Fig. 1A)**, potentially reflecting either the plasticity or small overall contribution of this hydrogen bonding interaction. Residue L452 – which is mutated to L452R in Delta and Kappa variants – is located within the periphery of the V_H_ F6 epitope and may contribute hydrophobic interactions with the V_H_ F6 residue F58 (**Fig. S3D)**. The peripheral nature of this interaction may explain the marginal sensitivity of V_H_ F6 binding to the L452R mutation (**Fig. 1A)**. In contrast, F490L and F490S mutations attenuate and completely abrogate V_H_ F6 binding respectively (**Fig. 1A)**, as can be rationalized by the location of F490 within both the FR and CDR3 binding interfaces (**Fig. S3E)**. The lack of significant interactions with VOC mutated residues provides a structural basis for the broad activity of F6.

The resolved F6/Beta spike structure may also explain the binding of V_H_ F6 to Omicron. According to the resolved F6/Beta RBD, 13 out of 15 omicron RBD mutations are located outside of the F6 epitope (**Fig. 2F)**, and the remaining two mutations, G446S and Q493R are in the peripheral region of F6 footprint. Importantly, our RBD mutants ELISA showed the G446S and Q493R mutations did not significantly disturb F6 binding (**Fig. 1A)**. Structure modeling and molecular dynamics (MD) simulations were performed to examine the interfacial interactions and showed that the complex formed between the Omicron variant RBD and F6 stably retained the same structural features as the cryo-EM resolved F6-Beta RBD complex in triplicate runs of 800 ns. The mutation sites Q493R and Q498R intermittently formed new compensating salt bridges. Simulations and binding energy calculations repeated for the complexes of F6 with Beta and Omicron variants led to respective K_D_ values of 12.2±3.1 nM and 15.5±3.3 nM, which is in line with the Blitz K_D_ (**Fig. S4**).

### Generation of a biparatopic antibody with enhanced neutralization of SARS-CoV-2 VOCs

To expand the V_H_ F6 epitope, with the aim of decreasing mutational escape, we designed a biparatopic Ab by combining V_H_ F6 and V_H_ ab8, which is a nAb with a distinct yet partially overlapping epitope compared to F6 (**Fig. S5A and S5B**). While ab8 is escaped by the Beta, Gamma, and Omicron variants **(Fig. S5C)**, ab8 is not escaped by the F490S and F490L mutations that escape V_H_ F6 binding (**Fig. S5D**). The biparatopic Ab was constructed by linking V_H_ F6 to V_H_ ab8 via a 5x(GGGGS) polypeptide linker with the C terminal fused to the human IgG1 fragment crystallizable region (Fc) (**Fig. 3A**). Addition of an Fc region has previously been demonstrated to both extend antibody serum half-life and activate the effector function of the immune system[48]. The biparatopic Ab, F6-ab8-Fc, broadly bound to SARS-CoV-2 VOC spike trimer proteins (**Fig. S5E**). Importantly, F6-ab8-Fc potently bound to the Omicron RBD with an EC_50_ of 19.1 nM as measured by ELISA and a *K*_*D*_ of 38.7 nM as measured by BlitZ (**Fig. 3B and 3C**). Furthermore, F6-ab8-Fc potently neutralized WT, Alpha, Beta, and Delta SARS-CoV-2 variants in both pseudovirus and live virus assays (**Fig. 3D and E**). F6-ab8-Fc neutralized Omicron variant psuedoviruses with an IC_50_ of 4.82 nM (**Fig. 3D and F**), which is significantly more potent than V_H_ F6 (IC_50_ =269 nM). Additionally, F6-ab8-Fc neutralization of other SARS-CoV-2 VOCs is also more potent than that of V_H_ F6 (**Fig. 3F**), prompting us to evaluate its *in vivo* viral inhibition.

**Fig. 3.**
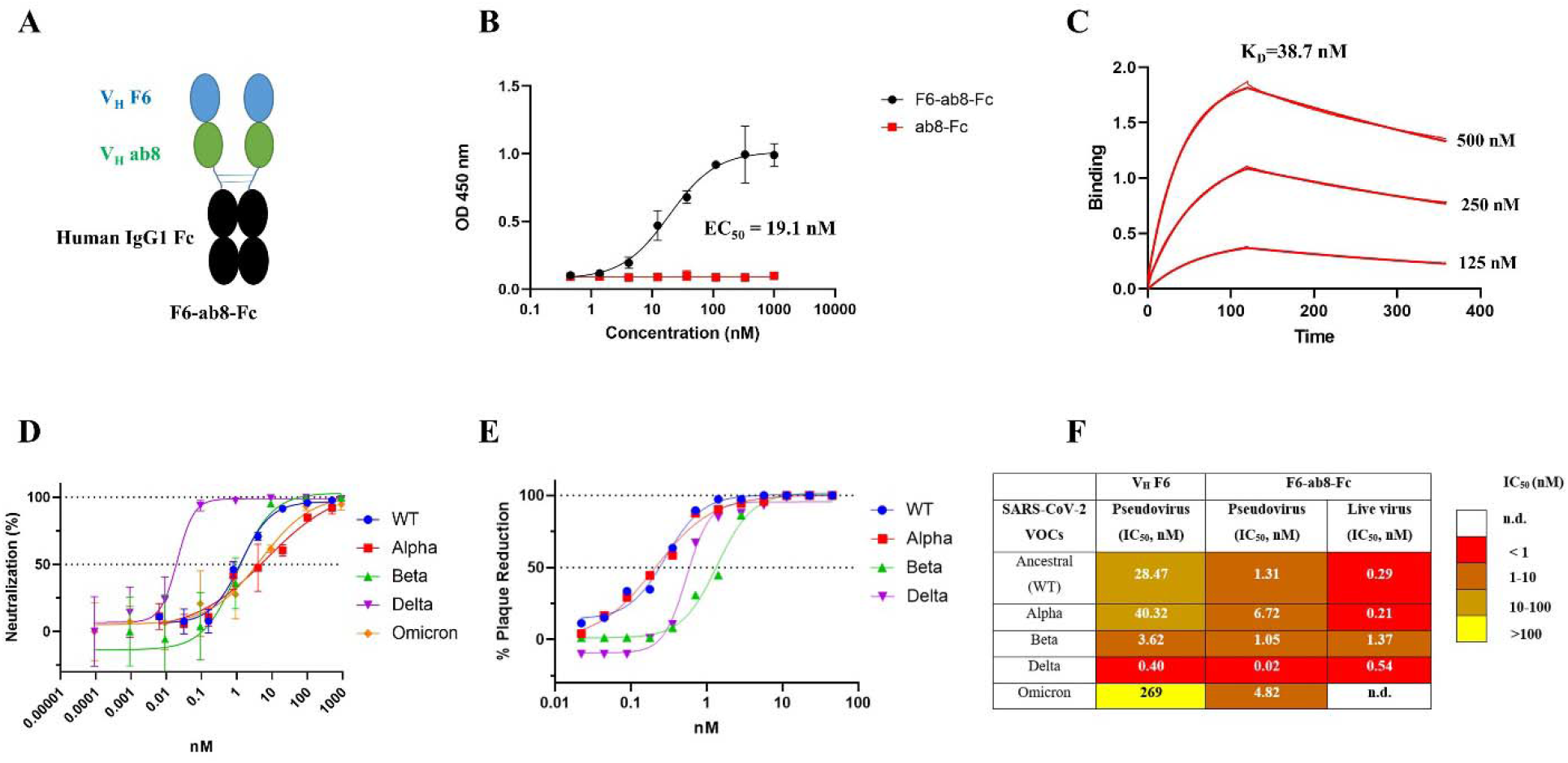
Construction of a biparatopic antibody (F6-ab8-Fc) that neutralizes various SARS-CoV-2 VOCs including Omicron. **A.** The scheme of the biparatopic antibody F6-ab8-Fc containing a tandem VH (F6-ab8) at the N terminal of the human IgG1 Fc. **B**. ELISA results of F6-ab8-Fc binding to the Omicron RBD protein (EC_50_= 19.1 nM). **C**. Binding kinetics of F6-ab8-Fc binding to the Omicron RBD tested by the BlitZ (K_D_=38.7 nM). **D-E**. Neutralization of SARS-CoV-2 WT, Alpha, Beta, and Delta variants pseudoviruses (**D**) and live viruses (**E**) by F6-ab8-Fc. **F**. Comparisons of virus neutralization IC50s by V_H_ F6 and F6-ab8-Fc.

### F6-ab8-Fc prophylactically and therapeutically reduces disease burden and protects from SARS-CoV-2 Beta variant mortality in mice

To evaluate the prophylactic and therapeutic efficiency of F6-ab8-Fc *in vivo*, we used a mouse-adapted SARS-CoV-2 infection model[51]. The Beta variant was chosen for *in vivo* protection experiments because it is relatively difficult to neutralize [36, 52]. Groups containing five mice each were administered a high dose of 800 μg or a low dose of 50 μg F6-ab8-Fc twelve hours pre-or twelve hours post-SARS-CoV-2 mouse-adapted 10 (MA10) Beta variant challenge. Mice were monitored for signs of clinical disease and viral titers in the lungs were measured four days after infection (**Fig. 4A**). Mice in the high-dose (800 μg) prophylaxis group were completely protected from mortality (0% morality). In contrast, 20% mortality was observed in the 800 μg therapeutic group and 40% mortality was observed in the 50 μg prophylactic group. 60% mortality was observed in the 50 μg therapeutic and control mAb group (**Fig. 4B**). Thus, F6-ab8-Fc can protect against mortality when given prophylactically at high doses. We observed greater than one log reduction in viral titer in the high-dose prophylactic and therapeutic groups after four days (**Fig. 4C**). Additionally, lung congestion scores, which is a gross pathologic score at the time of harvest, were lower in all four F6-ab8-Fc treated groups compared to the mAb control (**Fig. 4D**). Our results indicate that F6-ab8-Fc, is able to reduce lung viral replication *in vivo*, with prophylactic treatment being more effective than therapeutic treatment.

**Fig. 4.**
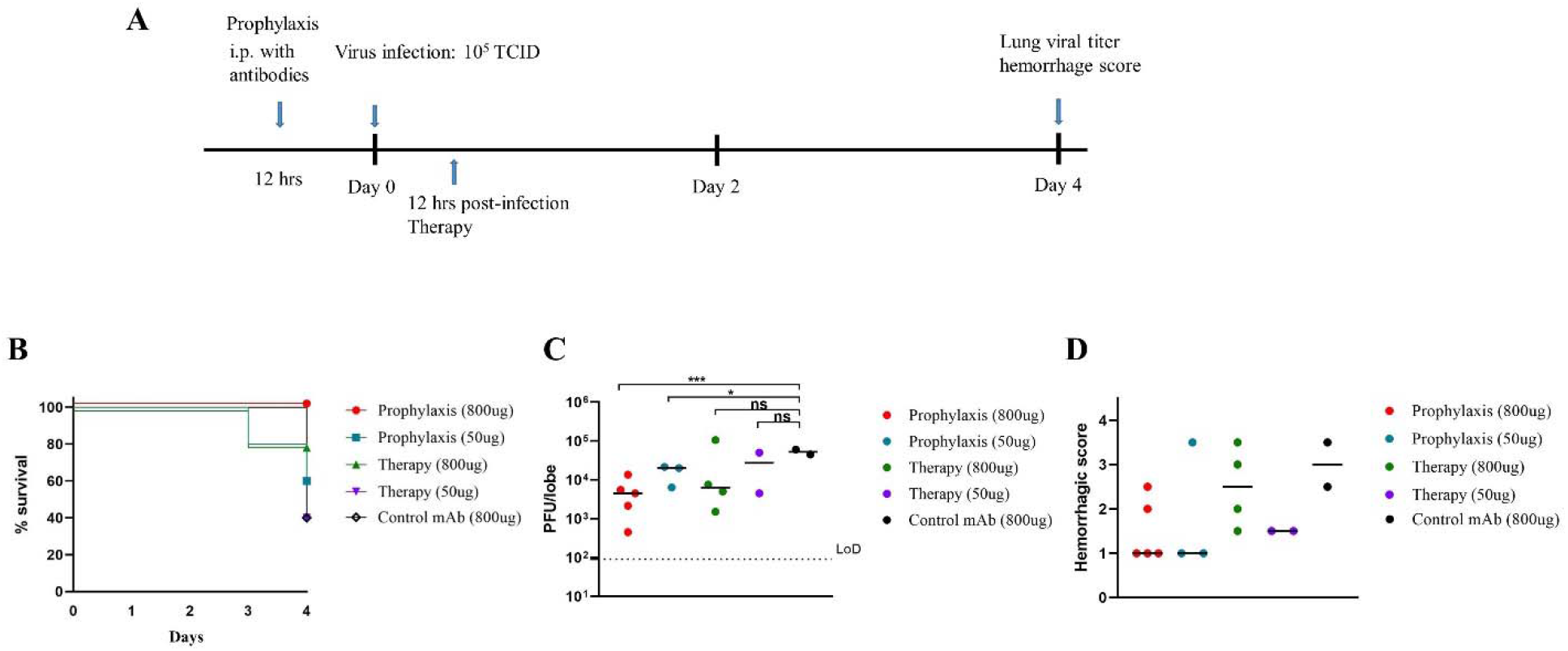
Evaluation of prophylactic and therapeutic efficacy of F6-b8-Fc in a mouse ACE2-adapted model. **A.** The overview of study design for evaluating F6-ab8-Fc efficacy in a SARS-CoV-2 mouse model. **B**. Percent survival curves for each F6-ab8-Fc treatment group as indicated. **C**. Viral titers (PFUs) in lung tissue for the F6-ab8-Fc treatment groups. The limit of detection (LoD) is 100 pfu/lobe. **D**. Lung hemorrhage scores of live mice. *T* tests were used to evaluate statistical differences. *p <0.05, **p <0.01, ***p < 0.001, ns. no significance.

## Discussion

The SARS-CoV-2 spike protein has accumulated numerous mutations that retain its ability to engage its receptor (hACE2), while evading neutralizing Abs [53]. The RBD is immunodominant and has accumulated several mutations that partially escape approved vaccines and the majority of clinical mAbs. A recent epitope binning and structural study classifies Ab epitopes across the RBD into six classes, with class 1-3 Abs targeting the top surface RBM region (thus competing with ACE2), and class 4/5 and class 6/7 Abs binding to the RBD outer and inner surfaces respectively [54]. Class 1-3 Abs are most likely to be rendered ineffective by K417N/T, E484K, and N501Y mutations which are found in Alpha, Beta, and Gamma VOCs. Currently, only a few RBM-targeting Abs are reported to neutralize the Omicron variant such as ACE2 mimicking Abs S2K146 [55] and XGv347 [56].

In this study, we developed a novel single domain (human V_H_) Ab, F6 that can broadly neutralize Alpha, Beta, Gamma, Delta, and Omicron variants. V_H_ F6 targets a class-4 epitope which spans the RBD peak and valley outer-face, and partially overlaps with the hACE2 binding interface. Importantly, the cryoEM structure of V_H_ F6 in complex with the Beta S protein revealed that VOC mutations lie either outside of the V_H_ F6 epitope (K417, E484, N501, N439) or within its periphery (L452, Q493, G446). The V_H_ F6 epitope bears high similarity to that of the full-length Ab A19-46.1, which can also neutralize the Omicron variant [57]. Unlike A19-46.1, V_H_ F6 is resistant to the L452R mutation, and can bind the RBD in both “up” and “down” conformations, probably due to its lower steric hindrance associating with its small size. The ability of an antibody fragment to bind both “up” and “down” RBD states is an attractive property given that the accessibility of its epitope is independent of RBD conformation [58]. Notably, V_H_ F6 adopts an uncommon angle of binding relative to the RBD, using its exposed FR regions and CDR3 to present a hydrophobic interaction interface. This interaction mode resembles that of llama/shark V_H_ Abs which use long CDR3s to fold against the FR2 region and collectively establish novel paratopes [59].

F6 primarily belongs to the class 4 Ab community, which also contains the highly potent and patient-derived Abs C002 [60] and A19-46.1 [57], and typically exhibits decreased binding to L452 and E484 mutated RBDs [61]. Additionally, the V_H_ F6 epitope partially overlaps with the with class 1-2 Abs which contain therapeutic Abs such as LY-CoV016 and REGN10933 [61] (**Fig. 2C**). The ability of the SARS-CoV-2 Omicron variant to escape class 1 and 2 Abs requires the development of Ab combinations (either cocktails or bi(multi)-specifics) targeting multiple epitopes. In this study, with the aim to target a broader epitope on the RBD, we generated a bispecific (biparatopic) Ab by combining F6 with the previously identified potent class-2 Ab domain V_H_ ab8 [48]. Although both Beta and Omicron variants were escaped by V_H_ Ab8, the biparatopic Ab, F6-ab8-Fc, potently neutralized all SARS-CoV-2 VOCs (including Omicron) *in vitro*. Importantly, V_H_ F6 also reduced lung viral titers in mice infected with the Beta variant and protected against mortality when administered prophylactically.

In summary, we identified a broadly neutralizing antibody domain (V_H_ F6) with a unique paratope and epitope, and which neutralized all SARS-CoV-2 VOCs. The F6 epitope may be targeted to elicit broadly neutralizing Abs and vaccines against circulating SARS-CoV-2 variants. The biparatopic Ab, F6-ab8-Fc, with its broad neutralization activity and *in vivo* activity presents a new Ab therapeutics against current SARS-CoV-2 VOCs.

## Materials and methods

### Antigen expression and phage panning

The SARS-CoV-2 RBD, S1 and S trimer mutants were ordered from Sino Biological (USA). The V_H_ F6 and V_H_ ab8 were expression in HB2151 bacteria cells as previously described[62, 63]. VH-Fc ab8, F6-ab8-Fc, and RBD-Fc were expressed with Expi293 cells as previously described[48, 62]. Expressed Protein purity was estimated as >95% by SDS-PAGE (Invitrogen) and protein concentration was measured spectrophotometrically (NanoVue, GE Healthcare). The panning process was described in detail in our previous protocol [64].

### ELISA

Ninety-six-well Elisa plates (Corning 3690) were coated with the RBD, S1 mutants or S trimer variants at a concentration of 5 μg/mL (diluted with 1xPBS) and incubated at 4 □ overnight (50 μL per well). The next day, plates were blocked with 150 μL 5% milk (Bio-RAD) in DPBS solution at room temperature for 2 hours. Primary antibodies were diluted with the same 5% milk blocking buffer and 1:10 or 1:3 serial dilution series were conducted, with 1 μM as the highest concentration. After 2 hours of blocking, the primary antibodies were added (50 μL per well) and incubated at room temperature for 2 hours. After 2 hours incubation, the plates were washed 4 times with 0.05% Tween 1xPBS (PBST) solution using a plate washer (BioTek). Secondary antibodies (anti-Flag-HRP or anti-Human Fc-HRP) were prepared with the same 5% milk at a dilution of 1:1000. 50 μL of secondary antibody was added into each well and incubated at room temperature for 1 hour. After 1 hour incubation, the plates were washed 5 times with PBST. 50 μL of TMB substrate (Sigma) was added into each well, allowed 1-2 minutes to develop color, then stopped with 50 μL H2SO4 (1M, Sigma) and the plate scanned at 450 nm absorbance. The ELISA results were analyzed using GraphPad Prism 9.0.2.

### BlitZ

Antibody affinities were measured by biolayer interferometry BLItz (ForteBio, Menlo Park, CA). For V_H_ F6 affinity determination, V_H_ F6 was biotinylated with BirA biotin-protein ligase standard reaction kit (BirA500, Avidity, USA). Streptavidin biosensors (ForteBio: 18–5019) were used for biotinylated V_H_ F6 immobilization. For F6-ab8-Fc affinity determination, Protein A biosensors (ForteBio: 18-5010) were used for immobilization. Dulbecco’s phosphate-buffered saline (DPBS) (pH = 7.4) was used for baseline and dissociation collection. The detection conditions used were: (I) baseline 30s; (II) loading 120 s; (III) baseline 30 s; (IV) association 120 s with a series of concentrations (1000 nM, 500 nM, 250 nM for V_H_ F6; 500 nM, 250 nM, 125 nM for F6-ab8-Fc); (V) dissociation 240 s. The K_a_ and K_d_ rates were measured by BLItz software and K_D_ was calculated for each antibody by the K_d_ /K_a_ ratio. For V_H_ F6 - V_H_ ab8 competition, Protein A biosensors (ForteBio: 18-5010) were used for RBD-Fc immobilization. The detection conditions used were (I) baseline 30s; (II) loading 120 s; (III) baseline 30 s; (IV) association 120 s with V_H_ ab8; (V) association 120 s with V_H_ F6.

### Electron Microscopy Sample Preparation and Data Collection

For cryo-EM, SARS-CoV-2 S trimer Beta mutant were deposited on grids at a final concentration of 2 mg/ml. Complexes were prepared by incubating S trimer Beta mutant with V_H_ F6 at a molar ratio of 1:10. Grids were cleaned with H2/O2 gas mixture for 15 s in PELCO easiGlow glow discharge unit (Ted Pella) and 1.8 μl of protein suspension was applied to the surface of the grid. Using a Vitrobot Mark IV (Thermo Fisher Scientific), the sample was applied to either Quantifoil Holey Carbon R1.2/1.3 copper 300 mesh grids or UltrAuFoil Holey Gold 300 mesh grids at a chamber temperature of 10°C with a relative humidity level of 100%, and then vitrified in liquid ethane after blotting for 12 s with a blot force of −10. All cryo-EM grids were screened using a 200-kV Glacios (Thermo Fisher Scientific) TEM equipped with a Falcon4 direct electron detector and data were collection on a 300-kV Titan Krios G4 (Thermo Fisher Scientific) TEM equipped with a Falcon4 direct electron detector in electron event registration (EER) mode. Movies were collected at 155,000× magnification (physical pixel size 0.5 Å) over a defocus range of −3 μm to −0.5 μm with a total dose of 40 e – /Å2 using EPU automated acquisition software (Thermo Fisher).

### Image Processing

A detailed workflow for the data processing is summarized in Supplementary Figure S2. All data processing was performed in cryoSPARC v.3.2 [65]. On-the-fly data pre-processing including patch mode motion correction (EER upsampling factor 1, EER number of fractions 40), patch mode CTF estimation, reference free particle picking, and particle extraction were carried out in cryoSPARC live. Next, particles were subjected to 2D classification (just for evaluation of the data quality) and 3 rounds of 3D heterogeneous classification. The global 3D refinement was performed with per particle CTF estimation and high-order aberration correction. Focused refinement was performed with a soft mask covering the down RBD and its bound V_H_ F6. Resolutions of both global and local refinements were determined according to the gold-standard FSC [66].

### Model Building and Refinement

Initial models either from published coordinates (PDB code 7MJI) or from homology modeling (V_H_ F6)[67] were docked into the focused refinement maps or global refinement maps using UCSF Chimera v.1.15[68]. Then, mutation and manual adjustment were performed with COOT v.0.9.3[69], followed by iterative rounds of refinement in COOT and Phenix v.1.19[70]. Model validation was performed using MolProbity[71]. Figures were prepared using UCSF Chimera, UCSF ChimeraX v.1.1.1[72], and PyMOL (v.2.2 Schrodinger, LLC).

### Molecular dynamics simulations of SARS-CoV-2 Omicron RBD complexed with F6, and evaluation of binding energies

We constructed a structural model for the Omicron RBD complexed with F6 using the cryo-EM structure of F6/Beta RBD complex as template, and constructed the system for molecular dynamics simulations of this complex using the CHARMM-GUI Solution Builder module[73]. The resolved N-linked glycans and disulphides were included in the model, along with explicit water molecules to cover a distance 10 Å away from protein edges. Sodium and chloride ions corresponding to 0.15 M NaCl were included. This resulted in a simulation box of 94×94×94 Å^3^. CHARMM36 force field with CMAP corrections was used for the protein, water, and glycan molecules[74, 75]. All MD simulations were performed using NAMD (version 2.13)[76] with the protocol adopted from earlier work[77]. Simulations were performed in triplicates with 100 ns each for the Omicron RBDs complexed with F6. Binding free energies ΔGb_inding_ were evaluated using PRODIGY[78], and binding dissociation constants, K_D_, using K_D_ = exp(ΔG_binding_/RT) × 10^9^ (in nM) with RT = 0.6 kcal/mol at T = 300K. ΔG_binding_ histograms were generated based on 800 snapshots evenly collected during the MD simulation time interval 20 < t ≤ 100 ns for each run.

### Pseudovirus Neutralization Assay

SARS-CoV-2 spike Wuhan-Hu-1 (+D614G), Alpha, Beta, Gamma, Delta, and Omicron protein genes were synthesized and inserted into pcDNA3.1 (GeneArt Gene Synthesis, Thermo Fisher Scientific). HEK293T cells (ATCC, cat#CRL-3216) were used to produce pseudotyped retroviral particles as described previously[79]. 60 hours post transfection, pseudoviruses were harvested and filtered with a 0.45 μm PES filter. HEK293T-ACE2-TMPRSS2 cells (BEI Resources cat# NR-55293) were seeded in 384-well plates at 20 000 cells for neutralization assays. 24 hours later, normalized amounts of pseudovirus preparations (Lenti-X™ GoStix™ Plus) were incubated with dilutions of the indicated antibodies or media alone for 1 h at 37°C prior to addition to cells and incubation for 48 h. Cells were lysed and luciferase activity assessed using the ONE-Glo™ EX Luciferase Assay System (Promega) according to the manufacturer’s specifications. Detection of relative luciferase units (RLUs) was measured using a Varioskan Lux plate reader (Thermo Fisher).

### Authentic SARS-CoV-2 Plaque Reduction Neutralization Assay

Neutralization assays were performed using Vero E6 cells (ATCC CRL-1586). One day before the assay, the Vero E6 cells (3 × 10^5^ cells) were seeded in 24-well tissue culture plates per well. Antibodies were serially diluted by two-fold with a starting concentration ranging from 4 μg/mL to 40 μg/mL (depending on the antibody being tested) and mixed with equal volume of 30-50 plaque forming units (pfu) of SARS-CoV-2. The following SARS-CoV-2 variants were used: isolate USA-WA1/2020 (NR-52281, BEI Resources); isolate hCoV-19/South Africa/KRISP-EC-K005321/2020 (NR-54008, BEI Resources); isolate hCoV-19/USA/CA_UCSD_5574/2020 (NR-54008, BEI Resources); and isolate hCoV-19/USA/PHC658/2021 (NR-55611, BEI Resources). The antibody-virus mixture was then incubated at 37°C in a 5% CO2 incubator for 1 hour before adding to the Vero E6 cell seeded monolayers. The experiments were performed in duplicate. Following 1 h incubation at 37 °C, an overlay media containing 1% agarose (2x Minimal Essential Medium, 7.5% bovine albumin serum, 10 mM HEPES, 100 μg/mL penicillin G and 100 U/mL streptomycin) was added into the monolayers. The plates were then incubated for 48-72 hours and then cells were fixed with formaldehyde for 2 hours. Following fixation, agar plugs were removed, and cells were stained with crystal violet. To precisely titrate the input virus, a viral back-titration was performed using culture medium as a replacement for the antibodies. To estimate the neutralizing capability of each antibody, IC50s were calculated by non-linear regression using the sigmoidal dose response equation in GraphPad Prism 9. All assays were performed in the University of Pittsburgh Regional Biocontainment Laboratory BSL-3 facility.

### Evaluation of F6-ab8-Fc Prophylactic and Therapeutic Efficacy with SARS-CoV-2 mouse Models

Eleven to twelve-month old female immunocompetent BALB/c mice (Envigo, stock# 047) were used for SARS-CoV-2 in vivo Prophylactic and Therapeutic experiments as described previously[51]. Each group contains five mice and five mice per cage (contain one mouse from each group) and fed standard chow diet. To evaluate the prophylactic efficacy of F6-ab8-Fc, mice were intraperitoneal (i.p.) injection with 800 μg or 50 μg of F6-ab8-Fc 12 hours prior virus infection. Mice were infected intranasally with 10^5^ plaque-forming units (PFU) of mouse-adapted SARS-CoV-2 B.1.351 MA10. For evaluating the therapeutic efficacy of F6-ab8-Fc, mice were intraperitoneal injection with 800 μg of or 50 μg of F6-ab8-Fc12 hours following infection. 4 days after virus infection, mice were sacrificed, and lungs were harvested for viral titer by plaque assays. The caudal lobe of the right lung was homogenized in PBS. The homogenate was 10-fold serial-diluted and inoculated with confluent monolayers of Vero E6 cells at 37°C, 5% CO_2_ for 1 hour. After incubation, 1 mL of a viscous overlay (1:1 2X DMEM and 1.2% methylcellulose) is added into each well. Plates are incubated for 4 days at 37°C, 5% CO2. Then, the plates are fixation, staining, washing and dried. Plaques of each plate are counted to determined virus titer. The study was carried out in accordance with the recommendations for care and use of animals by the Office of Laboratory Animal Welfare (OLAW), National Institutes of Health and the Institutional Anll e17 Cell 184, 4203–4219.e1–e18, August 5, 2021 Article imal Care. All mouse studies were performed at the University of North Carolina (Animal Welfare Assurance #A3410-01) using protocols (19-168) approved by the UNC Institutional Animal Care and Use Committee (IACUC) and all mouse studies were performed in a BSL3 facility at UNC.

## Supporting information

Supplementary Table S1, Figures and Figure Legends for S1 to S5

## Acknowledgments

We would like to thank the members of the Center for Antibody Therapeutics Dr. Jelev Dontcho, Yejin Kim, Du-San Baek and Xiaojie Chu for their helpful discussions. This work was supported by the University of Pittsburgh Medical Center. David R. Martinez is funded by a Hanna H. Gray Fellowship from the Howard Hughes Medical Institute and a Burroughs Wellcome Fund Postdoctoral Enrichment Program Award. RSB is supported by grants from the NIH AI132178 and AI108197. Work in the Subramaniam laboratory is supported by a Canada Excellence Research Chair Award and a grant from Genome BC, Canada. Ivet Bahar is funded by NIH awards P41GM103712 and R01GM139297. Simon M. Barratt-Boyes is funded by NIH R21AI47017.

## Author contributions

WL, DRM, SS, DSD, IB, RSB, JWM and SMB conceived and designed the research; WL and CC identified and characterized antibodies; WL made the V_H_ phage-display libraries. CC, AK, XL, CA, MGH and ZS characterized antibodies and made the stable cell lines. WC provided Omicron RBD proteins. AS and DRM performed the in vivo evaluation of inhibition of SARS-CoV-2 beta variant. JWS, DM, XZ and AMB resolved the cyroEM structure and did neutralization of pseudovirus. MDS, NE, KDM and JLJ performed the neutralization of SARS-CoV-2 pseudovirus. MMM, BRT, PMDSC tested the live virus neutralization. MHC, AB and IB carried out the molecular dynamic simulation of VH F6 binding to Omicron RBD. DSD, WL, CC, JWS, MHC and IB wrote the first draft of the article, and all authors discussed the results and contributed to the manuscript.

## Data Availability

The data that support this study are available from the corresponding author upon reasonable request. The atomic models and cryo-EM density maps for V_H_ F6/Beta S trimer complex have been deposited into the Protein Data Bank (PDB) (ID: XXXX) and Electron Microscopy Data Bank (EMDB) (ID: XXXX).

