## Supplementary Table S1, Figures and Figure Legends for S1 to S5 for "Potent Neutralization of Omicron and other SARS-CoV-2 Variants of Concern by Biparatopic Human VH Domains"

**This PDF file includes:**

Table S1, Figures and Figure Legends for S1 to S5

**Table S1. CryoEM density and model processing and validation parameters**

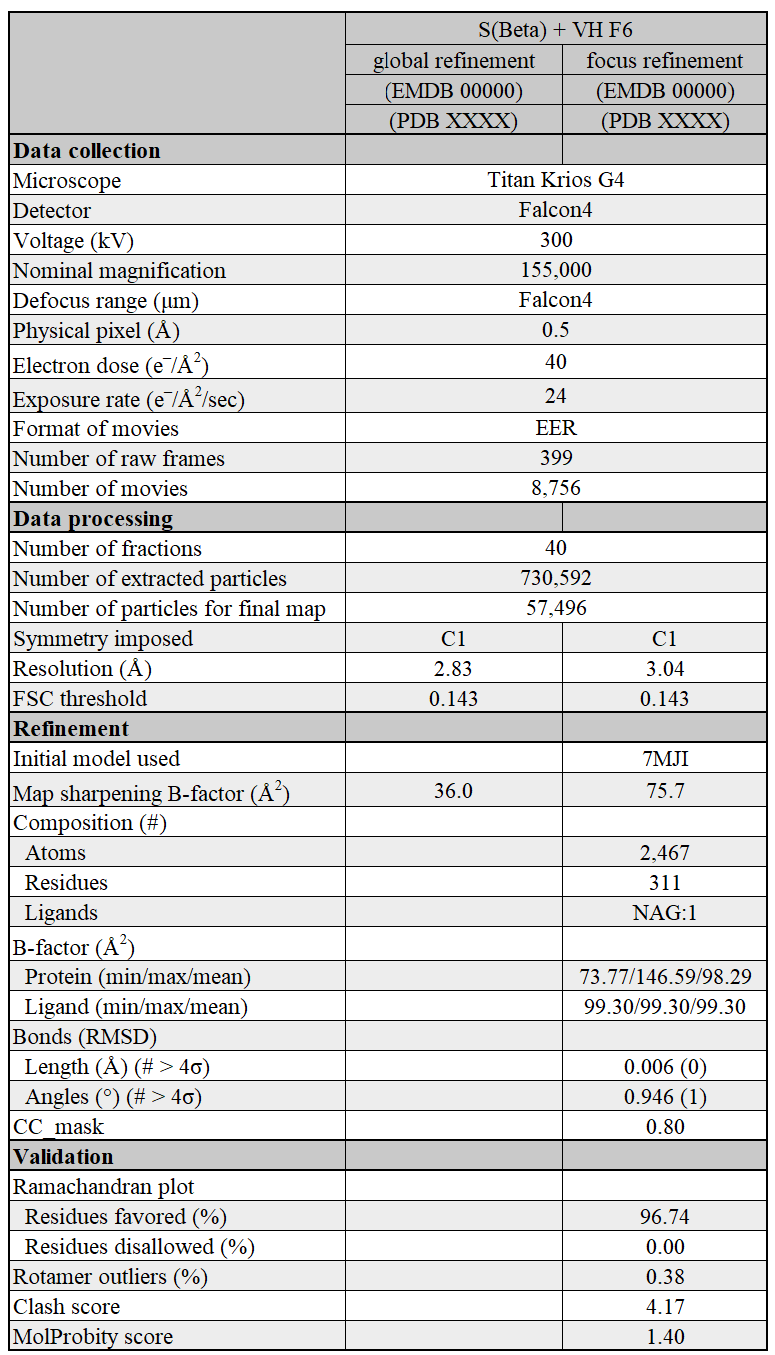

**Supplementary Figures and Figure Legends**

**Figure S1**

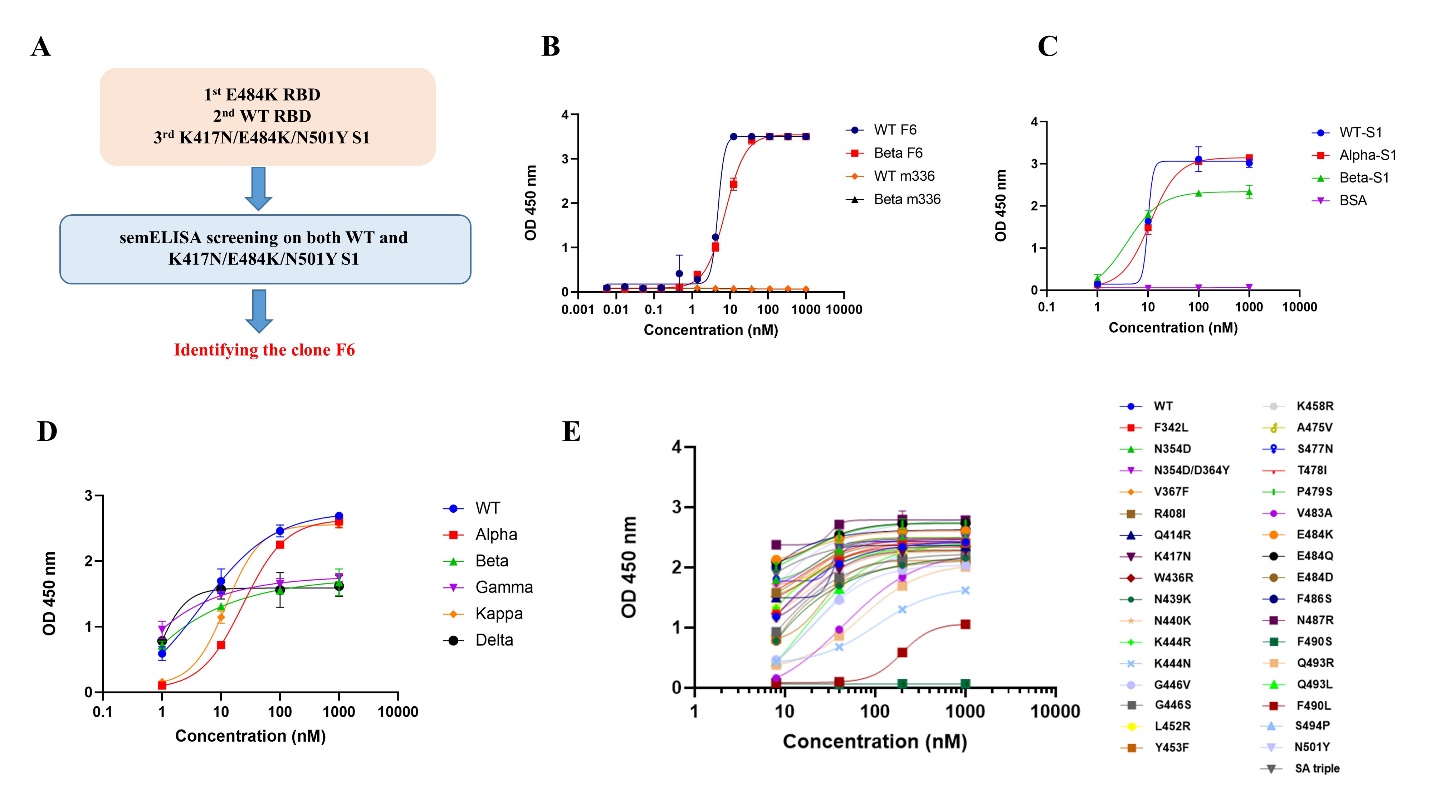

**Figure S1. Identification of V_H_ F6 by a sequential phage panning strategy, which broadly binds to SARS-CoV-2 VOCs spike and RBD proteins. A.** Overview of phage panning strategy. Three rounds of panning were performed using the RBD E484K mutant for the first round, WT RBD for the second round and K417N/E484K/N501Y S1 protein for the third round of panning followed by supernatant expression monoclonal (sem)ELISA screening. **B.** ELISA results of V_H_ F6 binding to the WT and Beta RBD proteins. The MERS-CoV antibody IgG1 m336 was used as a negative control. **C.** V_H_ F6 binding to the SARS-CoV-2 WT, Alpha and Beta S1 proteins. BSA was used as a negative control. **D.** V_H_ F6 binding to VOC S trimer proteins measured by ELISA. **E.** V_H_ F6 binding to naturally occurring RBD mutants. ELISA experiments were performed in duplicate and error bars denote ± SD, n=2.

**Figure S2**

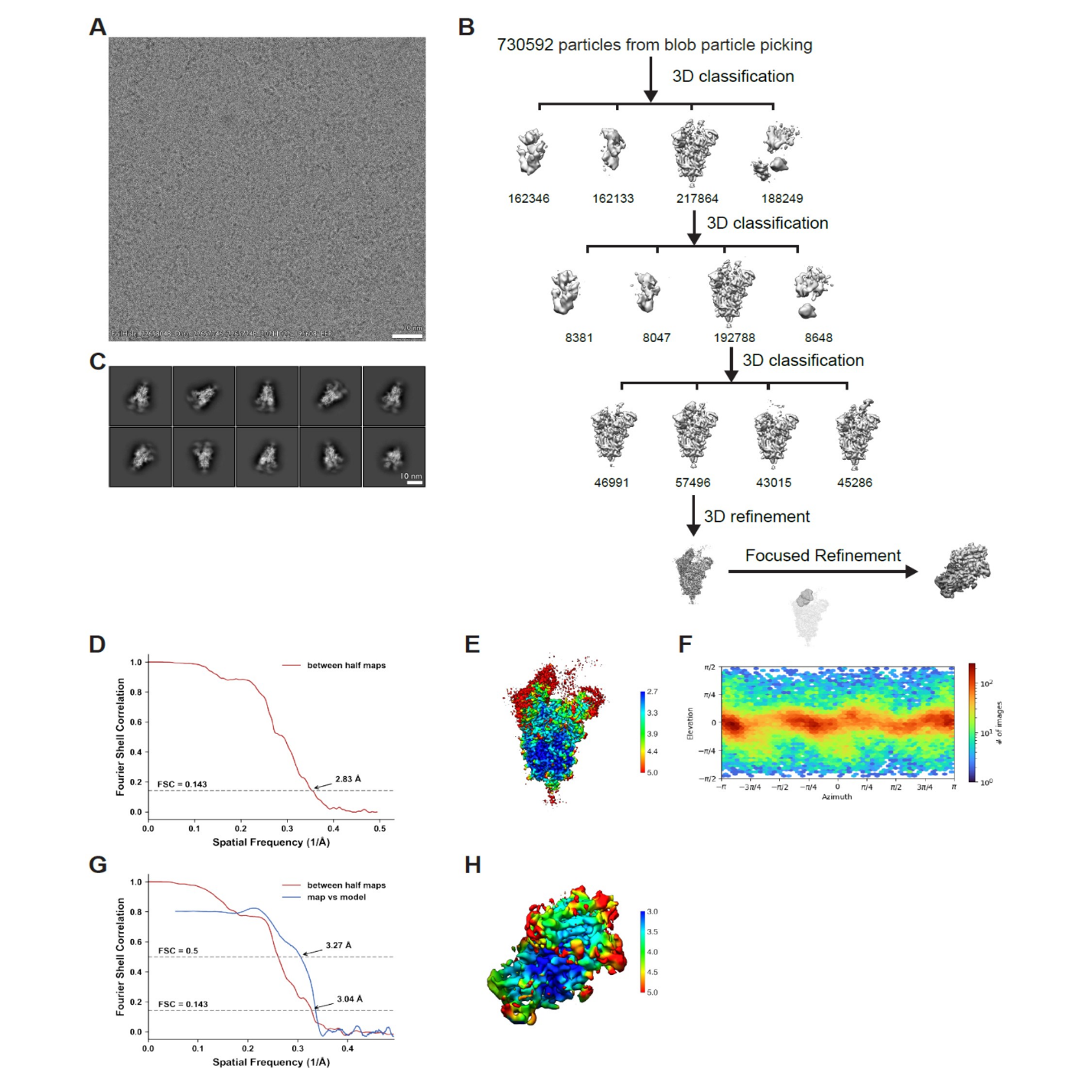

**Figure S2. Cryo-EM data processing and validation for the V_H_ F6 - Beta spike trimer complex.** **A.** Representative cryo-EM micrograph. **B.** Workflow of cryo-EM image processing. **C.** Representative 2D classes. **(D-F)** FSC curves (**D**), local resolution (**E**) and viewing direction distribution plot (**F**) of the global refinement. (**G-H**) FSC curves (**G**) and local resolution (**H**) of the focused refinement.

**Figure S3**

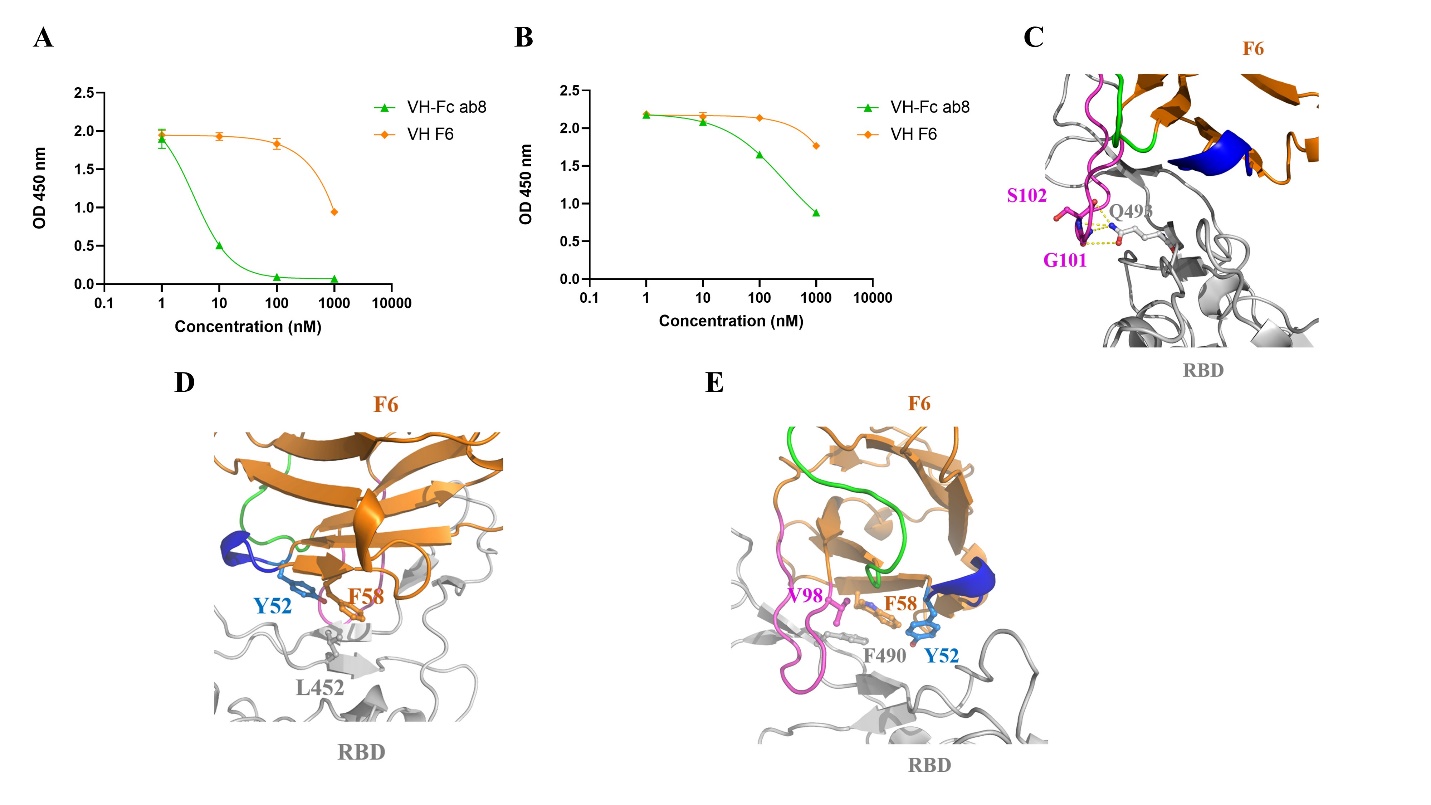

**Figure S3. ACE2 competition for V_H_ F6 and interfacial non-covalent interactions between F6 and RBD.** Competition ELISA of V_H_ F6 and V_H_-Fc ab8 with hACE2 for binding to the RBD (**A**) and S trimer (**B**). **C-E.** V_H_ F6 - RBD interaction interface focusing on residues Q439 (**C**), L452 (**D**) and F490 (**E**). The RBD is shown as a gray cartoon, and V_H_ F6 as an orange cartoon with CDR1, CDR2 and CD3 highlighted with green, blue, and magenta colors respectively.

**Figure S4**

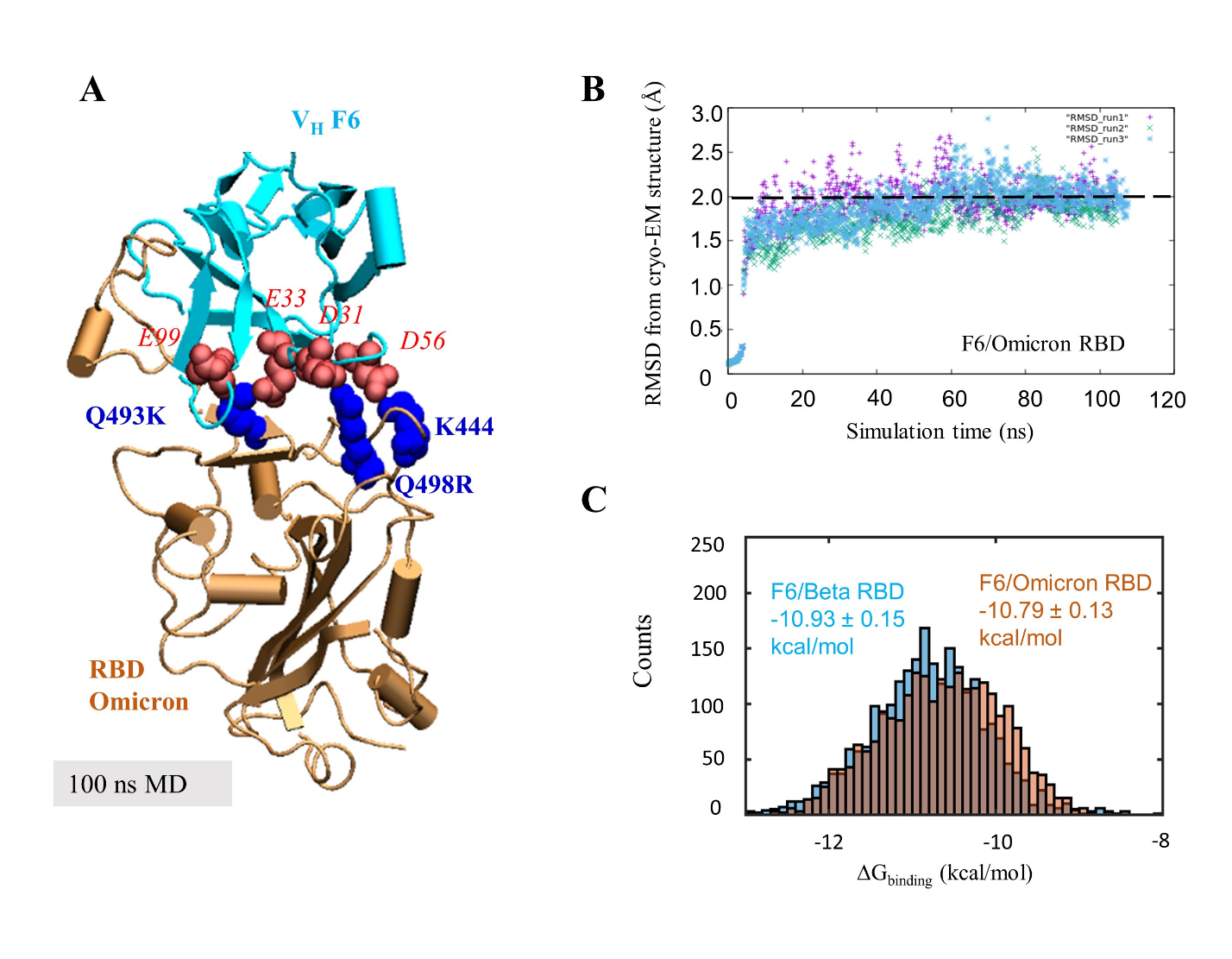

**Figure S4. Modeling of the structure, equilibrium dynamics, and binding energetics of the complex between VH F6 and Omicron RBD. (A) Structural model generated for the Omicron RBD-F6 complex**. The diagram shows the conformation stabilized after 100 ns MD simulations, which retains features comparable to those of the cryoEM resolved F6-Beta RBD complex. **(B) Stability of the F6/Omicron RBD complex near the conformation resolved for F6/Beta RBD complex.** The root-mean-square deviations (RMSDs) in the structural coordinates of F6/Omicron RBD with respect to the resolved F6/Beta RBD complex is shown as a function of simulation time. *Purple, blue and green* represent results taken from three different runs. The convergence to 2.0 ± 0.5 Å robustly reproduced in three different runs indicates the stability of the model that is closely similar to the cryo-EM structure resolved for F6/Beta RBD complex. (**C)** **Histogram of the binding energies**. Calculated based on three independent MD runs (800 evenly collected snapshots between 20 ns to 100 ns from each trajectory). The computed binding dissociation constants based on the average binding free energies are 12.2±3.1 and 15.5±3.3 nM, for the F6/Beta RBD and F6/Omicron RBD, respectively. Standard deviations are estimated based on three different runs.

**Figure S5**

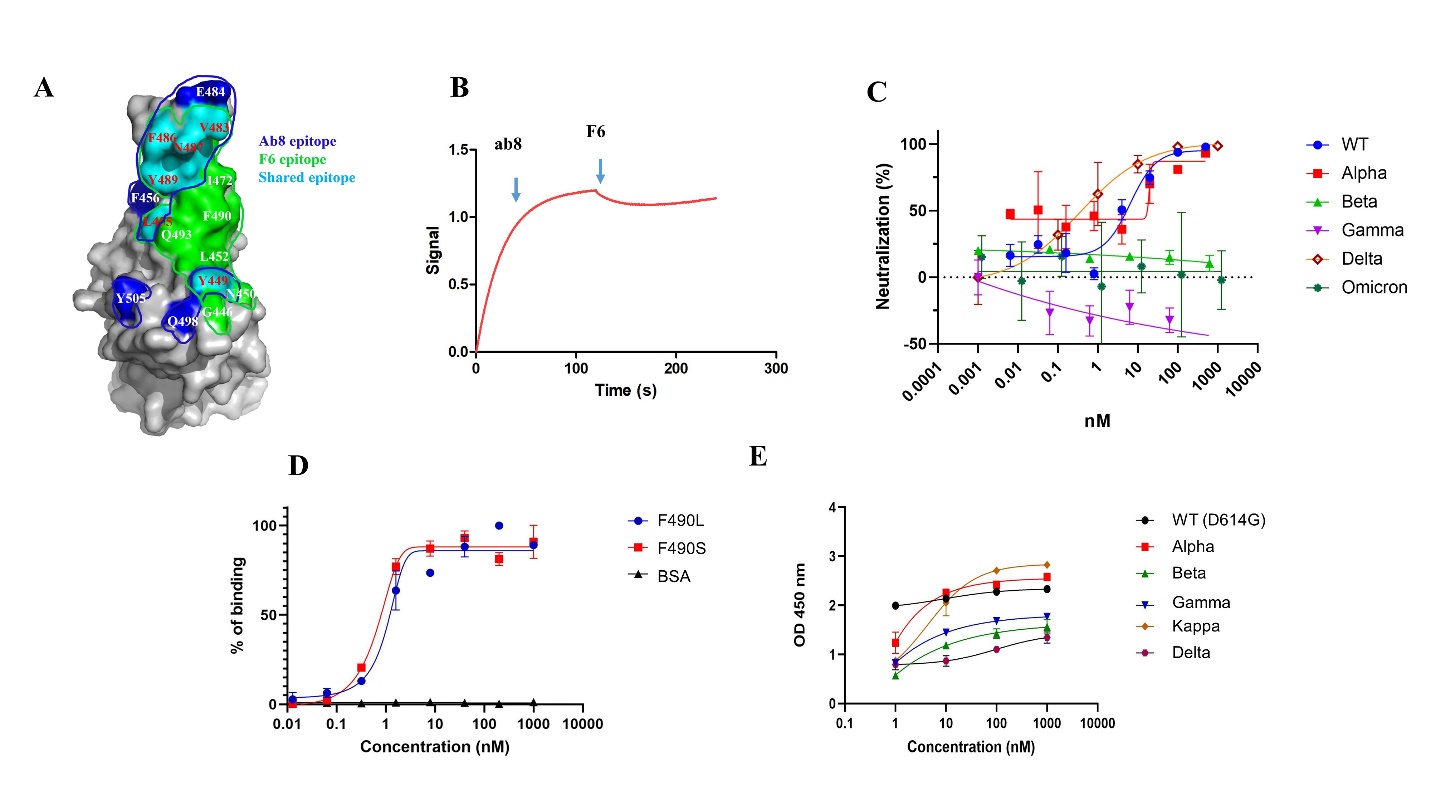

**Figure S5. Rational design of the biparatopic F6-ab8-Fc and its binding to SARS-CoV-2 VOCs proteins by ELISA. A.** Comparison of the F6 epitope (green) with the ab8 epitope (blue) on RBD surface. **B.** Competition of F6 with ab8 for binding to RBD as measured by BlitZ. **C.** Neutralization of SARS-CoV-2 VOC pseudoviruses by ab8, which is escaped the by the Beta, Gamma, and Omicron variants. Experiments were performed in triplicate and error bars denote ± SD, n=3. **D.** ab8 is able to bind to the SARS-CoV-2 RBD mutants, F490L and F490S, which escape V_H_ F6 binding. **E.** ELISA results of F6-ab8-Fc binding to VOC S trimer proteins. Experiments were performed in duplicate and error bars denote ± SD, n=2.
